# When the ostrich-algorithm fails: Blanking method affects spike train statistics

**DOI:** 10.1101/166611

**Authors:** Soheil Mottaghi, Kevin Joseph, Olaf Christ, Thomas J. Feuerstein, Ulrich G. Hofmann

## Abstract

Electrophysiological recordings of neuronal tissue face particular challenges when attempted during electrical stimulation, both in vivo and in vitro. Electrical stimulation may produce undesired electronic artifacts and thus render the recorded signal only partially useful. A commonly used remedy for these artifacts is to temporarily ground the input during the stimulation pulses. In the following study, we quantify the effects of this method on the spike train count, which is called "blanking". Starting a from theoretical standpoint, we deduce a loss of countable action potentials, depending on: width of the blanking window, Frequency of stimulation and neuronal activity. Calculations are corroborated by actual high SNR single cell recordings. We have to state, for therapeutically relevant frequencies of 130 Hz and realistic blanking windows of 2 ms, up to 27% of actual existing spikes are lost. We strongly advice careful and controlled use of blanking circuits when spike rate quantification is attempted.

## 3. Introduction

Direct electrical stimulation of neuronal tissue is used to provide insights into neuronal responses (Tehovnik 1996; Brindley & Lewin 1968; McCreery et al. 1990; Penfield 1958), to influence brain networks (Pezaris & Reid 2007; Xie et al. 2014; Choi et al. 2016) and to alleviate clinically relevant symptoms of neurological disorders (Kandel et al. 2000). To evoke a response, an electrical charge is delivered via electrodes to the region of interest, as described in seminal work by Rattay (Rattay 1989) and Tehovnik (Tehovnik et al. 2006). Unfortunately, while synchronously attempting the ‘read-out’ of neuron’s electrophysiological responses, the obligatory high level of amplification leads to signal contamination or even frequent saturation of the recording systems, i.e., the stimulation artifact (Hashimoto et al. 2002). The overlapping artifact may render the acquired neuronal signal difficult to analyze, which can be debilitating when neuronal spike statistics or responses are the focus of the investigation (Wagenaar & Potter 2002).

Stimulation artifact removal techniques within recorded data has been a thriving field, where linear and non-linear filtering (Gnadt et al. 2003; Whittington et al. 2005; Whittington et al. 2003; Parsa et al. 1998; Sennels et al. 1997) or artifact template subtraction utilizing a number of methods (Braack et al. 2013; Hashimoto et al. 2002; Zhiyue Lin & McCallum 1998) have been employed with mixed results.

Alternatively, several manufacturers of recording systems implement a hardware solution to prevent amplifier saturation by temporarily grounding its input with a so-called "blanking circuit". The input stage of the amplifier is thus protected from saturation and even damage, with a trigger signal that marks the stimulation pulse. The amplifiers are reconnected to the circuit after a user defined ‘blanking window’, usually following manufacturer’s recommendations. It is evident that during the ‘blanking window’, the signal values are set to zero (**Figure 1**), thus masking valuable neuronal signals.

**Figure 1:**
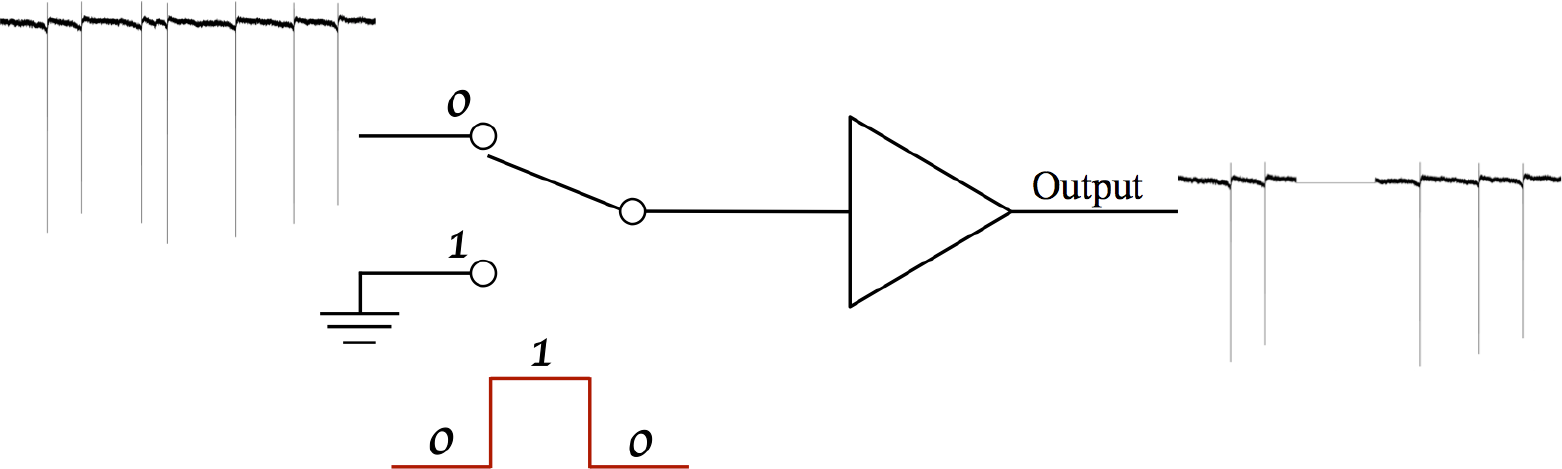
*Schematic of blanking circuit to prevent the amplifier from saturation in the presence of stimulation. The amplifier inputs are transiently grounded by the stimulation’s trigger for a time period defined by the respective user*.

This protective ‘blanking window’ needs to start latest upon onset of the actual stimulus pulse and needs to cover the complete discharge period the surrogate circuit of both the probe and electronics. This helps avoid cross talk between the stimulation and recording electrodes. This ‘artifact free’ signal is then further analyzed for changes in LFP or spike content (Yi et al. 2013; Qiu et al. 2015; Multichannel_Systems 2015; Heffer & Fallon 2008; Braack et al. 2013).

However, we caution users to exercise extreme care and consideration when using this seemingly bullet-proof method, as the chosen settings may very well have an influence on spike quantifications and indiscriminate use may introduce significant errors in real world recordings and spike counts.

There is a likelihood that the events of interest overlap with the blanking window, which results in event loss and potential consequences for subsequent neurobiological conclusions, requiring additional means to validate experiments (Xie et al. 2014). The following study investigates the influence of this widely used technique both theoretically and experimentally.

## 4. Theoretical evaluation - Probability of data loss

To assess the potential impact of periodic blanking on electrophysiological results based on spike statistics and counts, we utilize the spike probability of cells under investigation based on their interspike-interval-distribution. As previously described, spiking activity of neurons can be by a Poisson distribution and the interspike-interval-distribution (ISI) can be modeled by a Gamma distribution (Bair et al.1994). Cellular properties then define the interspike-interval-distribution and thus variability is to be expected between consecutive spikes.

The stimulation pulse triggered blanking circuit is designed to not just mask the actual pulse artifact with a width of *PW* and a repetition frequency *F_ST_*, but needs to include an additional safety margin to outlast the capacitive discharge, which could originate from either the system design or the electrode-tissue coupling. Setting the stimulation pulse width *PW*, a safety margin *T_M_* and the stimulation’s repetition frequency *F_ST_*, we seek to estimate the likelihood of a spike to fall within the blanking window *T_B_ =(PW+T_M_)*, where it will be lost.

The likelihood of a spike falling within TB is determined by the actual spiking probability of the neuron type under investigation, which is modeled by the Gamma distribution **Equation 1**, defined with scale and shape parameters, a and b respectively. The Gamma function Γ(a) and the mean value of the distribution μ are defined using **Equation 2** and **Equation 3**.

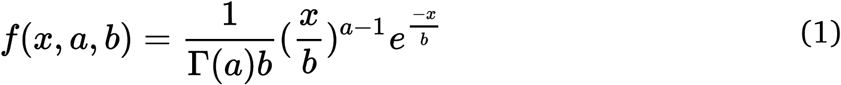

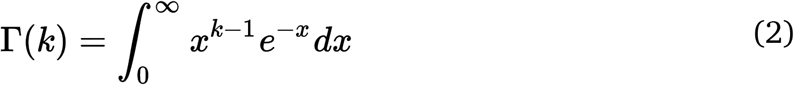

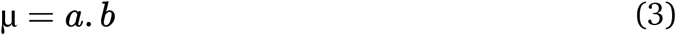

The cumulative probability of the Gamma distribution between x_1_ and x_2_ can be calculated using **Equation 4** and **Equation 5**. The maximal number of blanked bins N_b is determined using **Equation 6**, which integrates the ISI distribution of up to 99.95% of the counted events. If the number of blanked regions multiplied by the blanking window is larger than this x99.95 count, no spikes will be detected, as in **Equation 7**. The maximum probability of detection of non-blanked events is given by **Equation 8** and **Equation 9**. **Figure 2** illustrates blanking windows within the fitted Gamma distribution.

**Figure 2:**
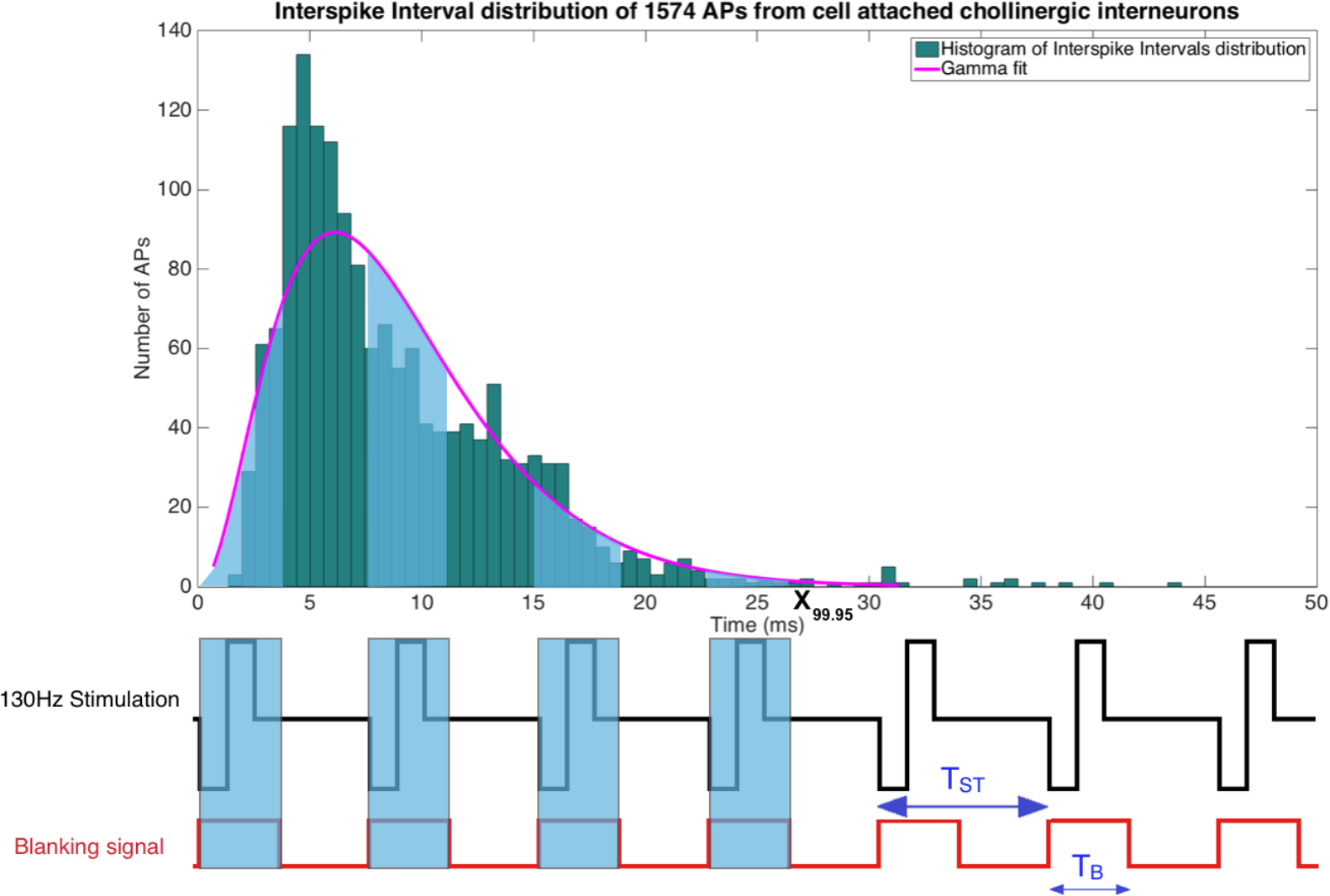
*Illustration of a periodic rectangular stimulation pattern with a repetition period T_ST._ The lower lines shows a sketch of a stimulation pulse train and the corresponding blanking signal, depicting its temporal extension beyond the actual stimulation pulse. Blanking windows are meant to suppress electrical signals thought to endanger sensitive amplifiers*.

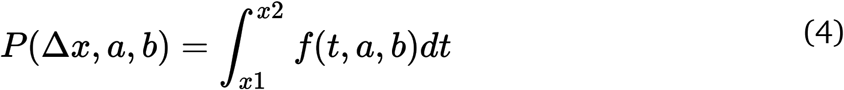

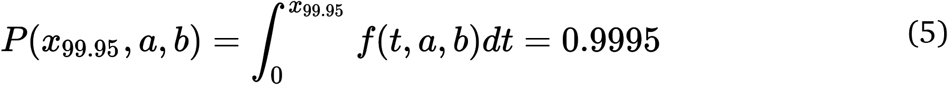

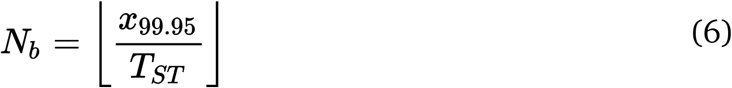

then

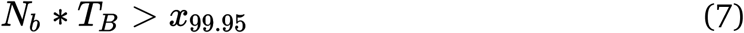

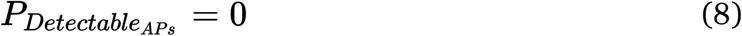

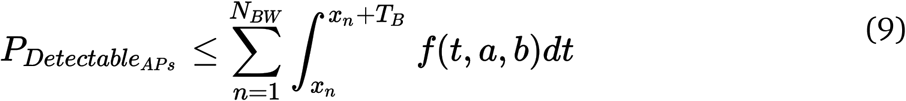

## 5. Experimental validation - Actual data loss

To test this theoretical approach, we performed cell-attached patch clamp recordings of cholinergic interneurons from the nucleus accumbens of rats in a slice preparation (**Figure 3**). This provides high SNR recordings with action potentials easily identified, despite non-saturating stimulus artifacts (**Figure 4**) providing ground truth measurements for spike detection and counts. Under these conditions, the stimulation artifact is clearly visible without obscuring the events, establishing a well-controlled test bed for the quantification of the blanking artifact.

**Figure 3:**
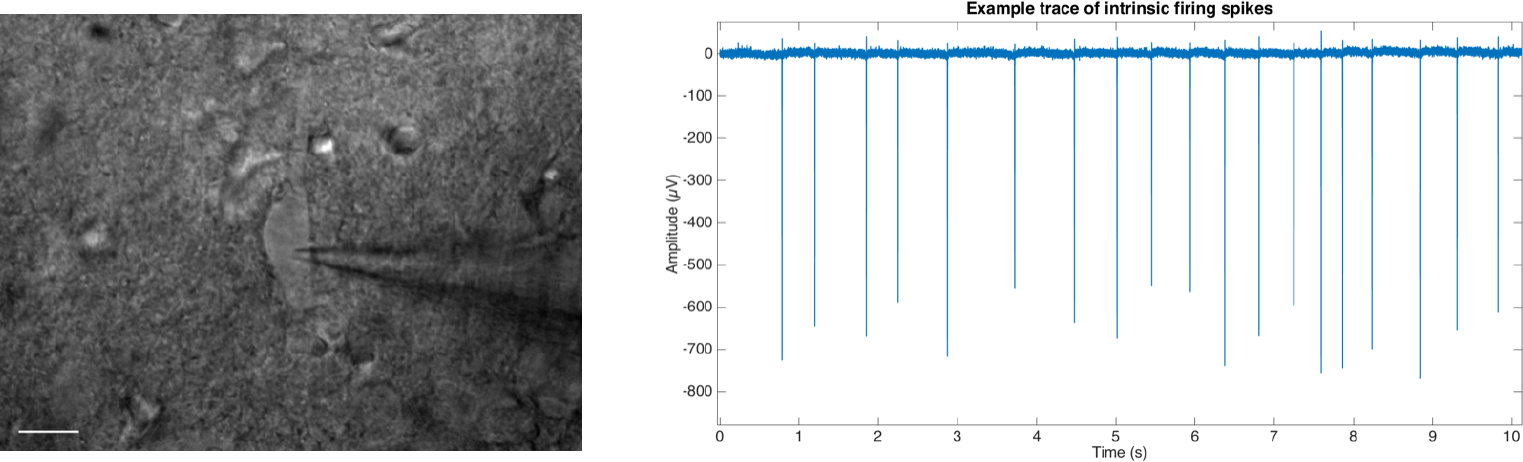
*(left) IR micrograph of a patched cholinergic interneuron. Scale bar is 20µm. (right) Example trace of intrinsic firing spikes under the depicted cell-attached patch clamp recording*.

**Figure 4:**
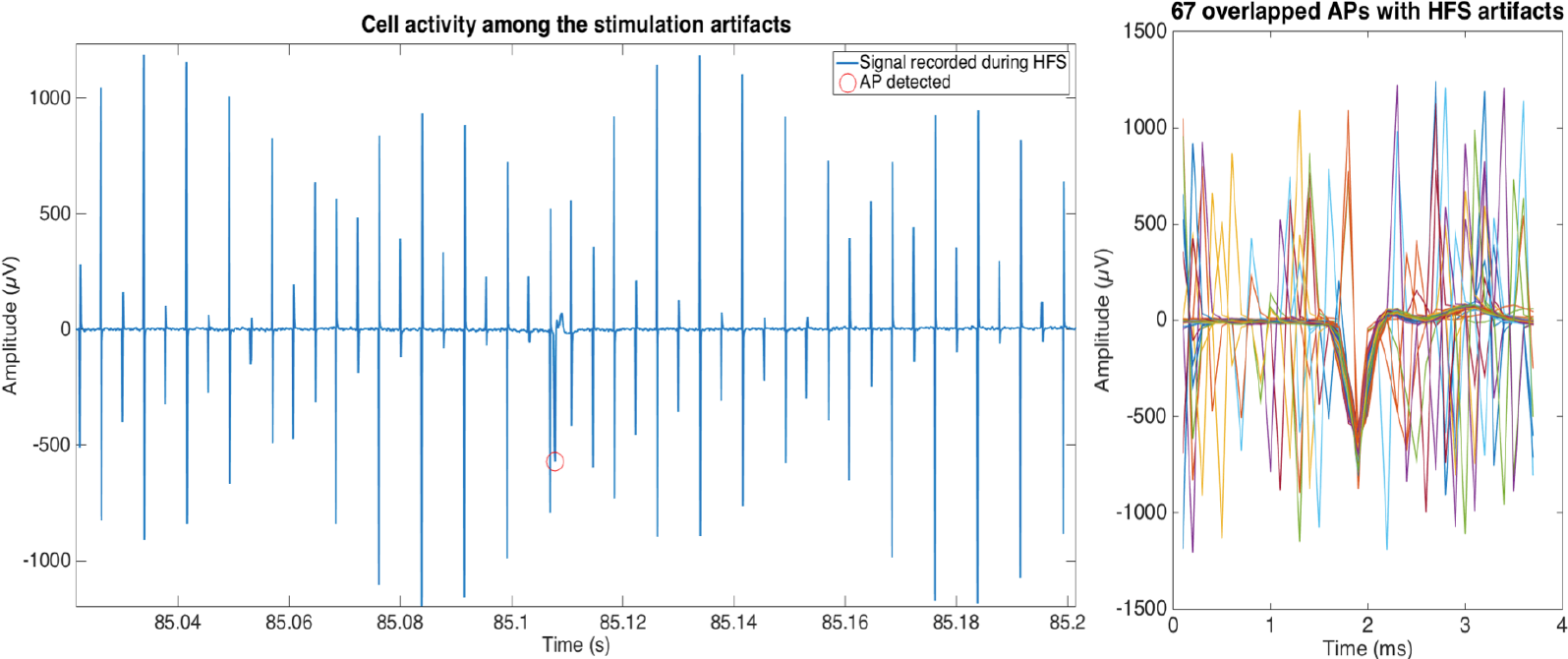
*(left) Example trace of stimulation artifact appearing in the cell-attached recording of a single cell. Stimulation parameters were biphasic, 130Hz, 12µA, 65µs pw. (right) Artistic rendering of possible occurrences of stimulation artifacts, in themselves not able to veil the prominent spike. Note: A blanking circuit is triggered by the occurrence of a stimulus pulse and will thus - depending on the timing-obscure the spike*.

### 5.1 Animals and tissue preparation

All protocols were approved by the responsible Animal Care Committee of the Regierungspräsidium Freiburg (permit X16/02A), and all efforts were made to minimize the number of animals used, with respect to statistical constraints. Male Wistar rats (3-4 weeks old) (Janvier, France), housed in groups of 4, under standard lighting (12h light-dark cycle), 22°C and 40% humidity were used in this study, and were allowed access to food and water *ad libitum*.

The rats were decapitated after an overdose of inhalation anesthetic (Forene, Baxter, USA), and the brains were quickly extracted and submerged in oxygenated ice-cold sucrose aCSF (artificial cerebrospinal fluid). Slices (300µm) were prepared using a vibratome (VT 1200, Leica, Germany) containing the nucleus accumbens with coordinates previously described (Varatharajan et al. 2015). After incubation at 37°C for 30 min in sucrose fortified aCSF, and a resting phase of 20 min at room temperature, the slices were ready for patch clamp experiments. All experiments were carried out at 32°C, controlled by an inline heater (Multi Channel Systems, Reutlingen, Germany) (Stuart et al. 1993). Neurons were visualized using infrared differential interference contrast (IR-DIC) video microscopy (XM-10, Olympus Corporation, Germany). Tissue slices were perfused with oxygenated aCSF containing: (in mM) 125 NaCl, 25 NaHCO3, 2.5 KCl, 1.25 NaH2PO4, 1 MgCl2, 2CaCl2, and 25 Glucose (pH 7.4 with 5% CO2).

### 5.2 Electrophysiological recordings

Recordings were carried out using a Multiclamp 700B Amplifier (Molecular Devices, USA) and the data was digitized using a CED 1401 Mark II (Cambridge Electronic devices, UK), acquired using a custom routine in IGOR Pro (Wavemetrics Inc., USA). Cholinergic interneurons from the Nucleus accumbens, were chosen for their tonic firing behavior (Xie et al. 2010).

Slices were submerged in the recording chamber and continuously perfused (4 ml/min) with oxygenated aCSF (34°C). Visualized patch-clamp recordings from the Nucleus accumbens were performed at 40× using infrared oblique-illumination (Olympus XM10, Olympus Corp., Japan) (**Figure 3**). Recording electrodes were pulled from thick walled borosilicate glass capillaries (2.0 mm O. D. and 1.2 mm I.D), filled with 150 mM NaCl (3-5 M) (Perkins 2006). To prevent clogging of the electrode tip while approaching targeted cells, a positive pressure was applied (40 mbar). Once a dimple was observed on the cell membrane surface, the pressure was released and a GΩ seal was seen to form almost instantly. If not, light suction pulses were applied until the seal was obtained. To prevent artificial depolarization of the cell membrane, the voltage was clamped such that a 0µA current injection was maintained. Action potentials could be observed almost instantaneously, a feature to be expected from tonically firing Cholinergic interneurons (**Figure 3**).

### 5.3 Electrical stimulation

Bipolar platinum-iridium (Pt/Ir) electrodes (GBCBG30, FHC Inc., Maine) with a central cathode (diameter 75µm, area 0.0062 mm^2^) and a concentric anode were used for stimulation within an average distance of 500 µm from the attached cell’s soma. The stimulation was carried out either using a commercially available PlexStim, (Plexon Inc, USA) or a stimulator developed in-house (Mottaghi et al. 2015) used to generate biphasic electrical stimuli with an amplitude of 12µA.

Stimulation parameters were chosen based on therapeutic HFS (High Frequency Stimulation) as described previously and. Both stimulator systems were used in charge-balanced mode and the pulse width was controlled on-line with an oscilloscope (TPS2012, Tektronix, Switzerland) in parallel to the electrode. Stimulation runs of 100 sec each were performed with a constant stimulation current of 12µA. Stimulation produces a visible artifact in the cell-attached extracellular recordings, despite the presumable charge balanced stimulation (**Figure 4**).

### 5.4 Experimental paradigms

Experimental runs were divided into phases of 100 sec each. Spike detection and counting was carried out offline in Matlab (Mathworks, USA) by an algorithm taking spike shapes into account. The algorithm is based on the detection of three main features of recorded events i.e. amplitude, rise-fall slopes and the peak shape respectively, defined during baseline recording. Based on arbitrarily chosen, overlapping signal windows, the algorithm detects potential events of interest despite artifact contamination by using the previously defined parameters. It then counts and archives the detected events for further statistical analysis (Appendix A). After each detection step, the event count was quantified and confirmed by a blinded expert to be compared against the detected events. **Figure 4** illustrates a detected event during the high frequency stimulation phase and an overlay of 67 more events to visualize possible artifact positions relative to them. After baseline recordings (100 s), the cells were stimulated for 100 s and then allowed to rest for 300 s before further experiments were carried out. The same cycle was repeated three times on each cell.

## 6. Results

The cell attached recording technique was chosen to obtain the best signal-to-noise ratio for action potentials despite ongoing electrical stimulation, yet did not protect against contamination. Even charge-balanced stimulation pulses frequently overlap action potentials with a substantial amplitude (**Figure 4**).

Signal deterioration, however, is worsened by setting all signal values to zero for the duration of the blanking window T_B_, as can be seen with an exemplary spike in **Figure 5**. It illustrates the successful removal of the stimulation artifacts at the cost of a clear change in event parameters due to the inconsiderate use of the blanking mechanism. Such distorted events are generally not detected by baseline-oriented methods like the one utilized here.

**Figure 5:**
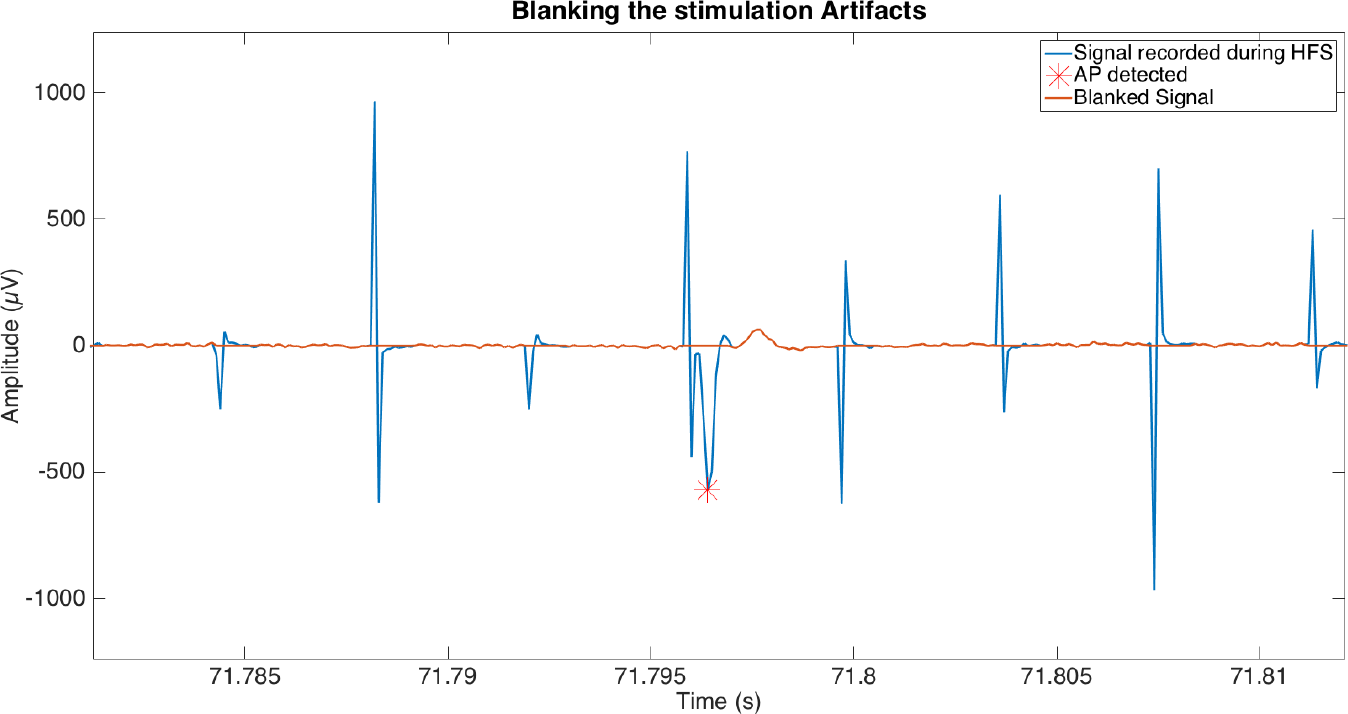
*Stimulation contaminated cell attached spike recording. Blanking of 1.5 ms very effectively removes all the stimulation artifacts but obviously shrouds parts of the spike as well (red line). Automated spike detection was then ineffective to detect the partially clipped spike in the blanked signal*.

As the above theoretical considerations illustrate, the usage of a blanking window will result in event loss, as given by **Equation 9**, **Figure 6**. The event loss for different blanking window widths is dependent on the stimulation/blanking frequency (**Figure 6**). The most commonly used blanking window of 2ms leads to a spike loss rate of up to 30 %, at therapeutically relevant frequencies. Wider blanking windows, motivated by sensitive amplifiers with longer saturation periods, clearly aggravates the situation and event loss may reach up to 80%.

**Figure 6:**
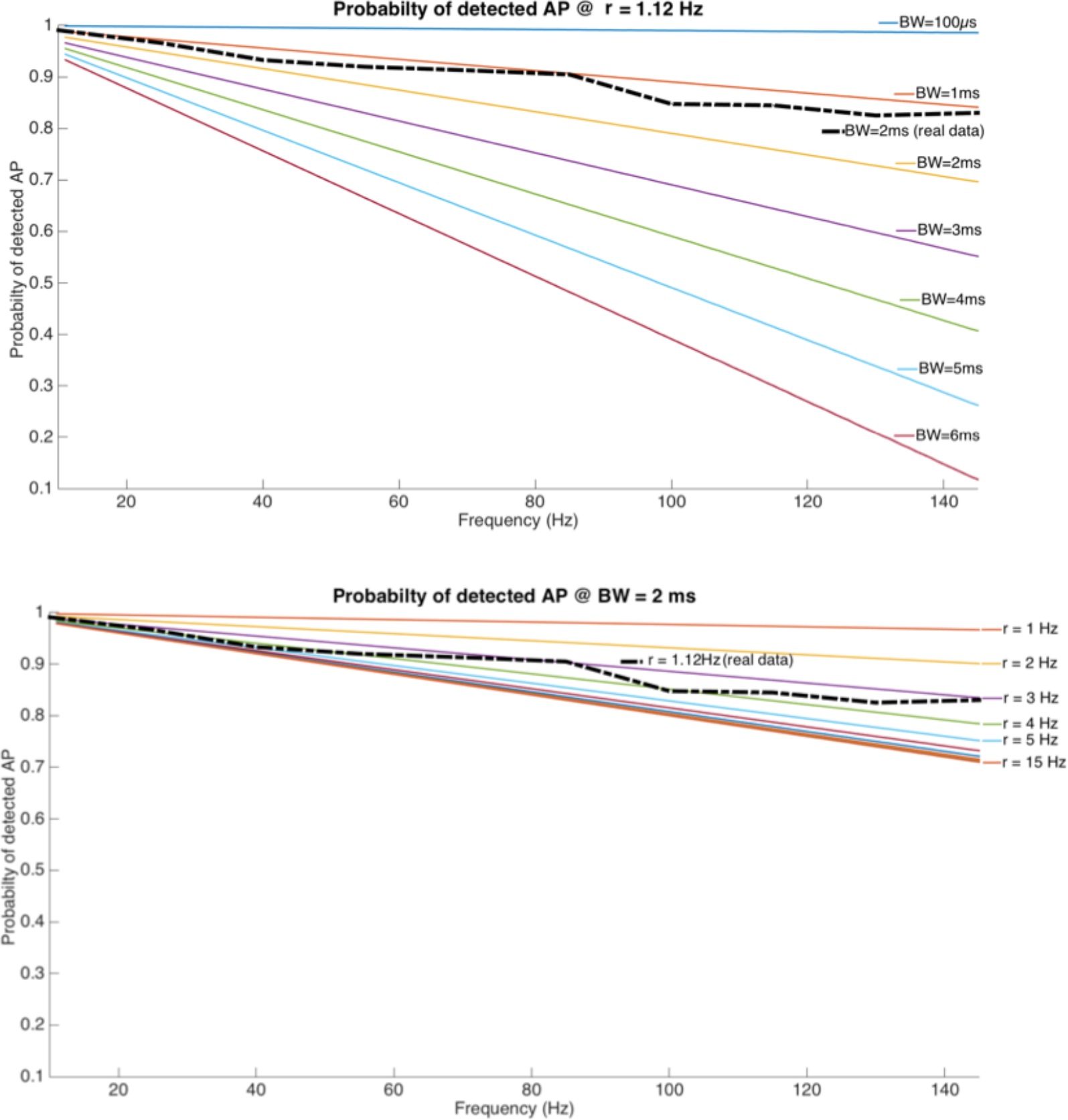
*(Top) Loss of spike count for a fixed spiking rate (r = 1.12 Hz) and stimulation pulse width p_w_, but increasing blanking window length and stimulation frequency. (bottom) Loss of spike count for a reasonable blanking window (T_b_=2msec) and increasing stimulation frequency and spike rate*.

Any potential loss in event counts is influenced by electrophysiological properties of the cells under investigation, given by their firing rates (**Figure 6**). Although intuitive, the percentage of lost events is directly proportional to the spike rate. However, for cell types with a spike rate> 3 Hz, the loss is independent of the spike rate and is influenced by the stimulation frequency alone. **Figure 7** top illustrates the inter spike interval (ISI) distribution of 1574 spikes from visually identified Cholinergic interneurons recorded in a cell attached configuration. The average firing rate ‘r’ was calculated as a product of its Gamma fit (Equation 1) parameters a=3.2051and b=277.6554 and resulted in r = 1.12Hz. The effect of a therapeutically relevant 130 Hz stimulation with a 2ms blanking window is demonstrated in Figure 7 bottom. Red bars represent histogram bins suppressed and thus lost from analysis by the blanking window. This is reflected in the experimental loss line (**Figure 6**).

**Figure 7:**
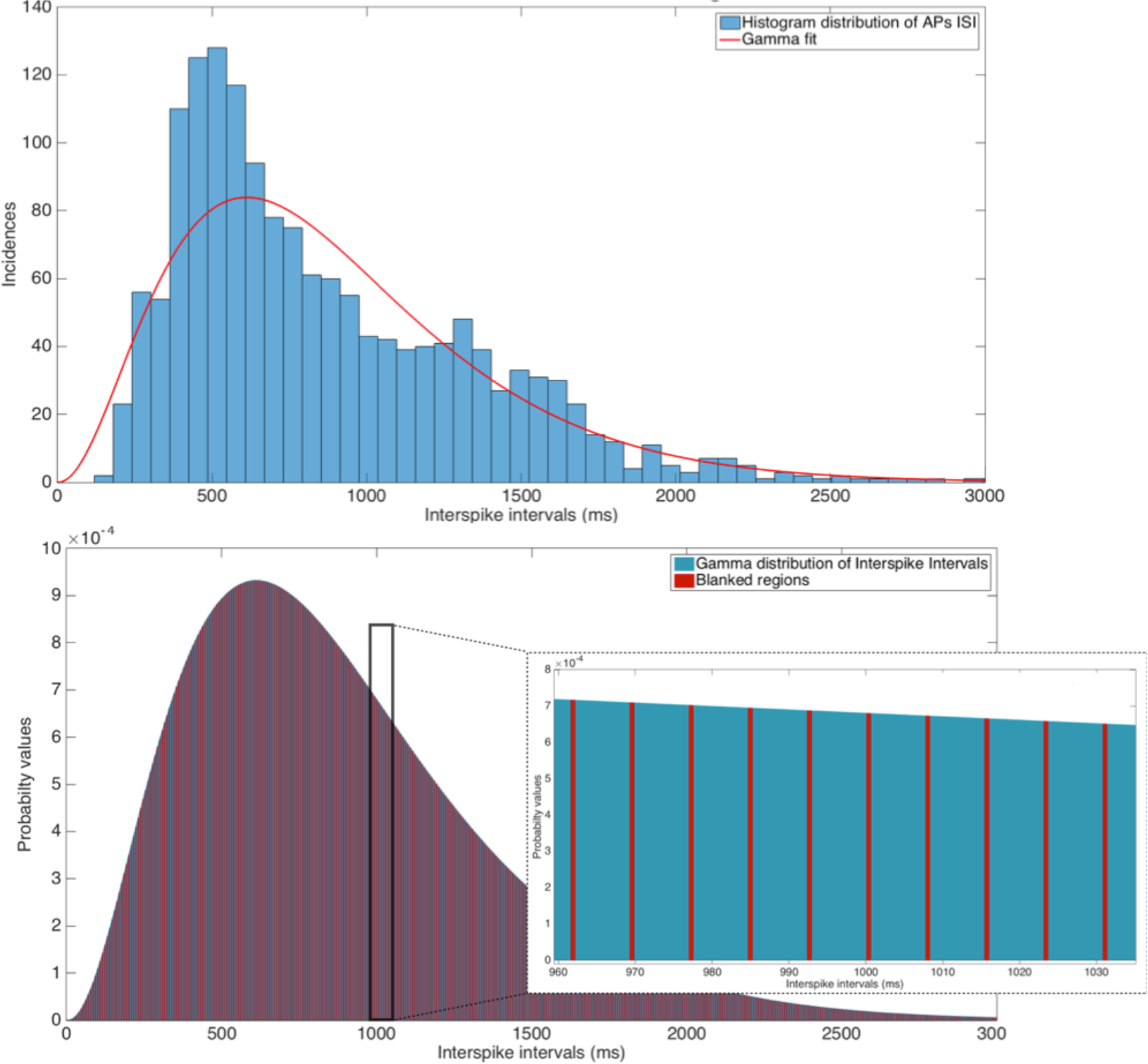
*(Top) Histogram of inter-spike intervals from 1574 cell attached cholinergic interneurons fitted by a Gamma distribution with a=3.2051 and b=277.6554. (Below). Illustration of a 2 ms blanking window suppressing a 130 Hz stimulation. Insert shows a zoom into the ISI period between 960 and 1035 ms. Scale of ordinate is equal for the ISI distribution and the insert*.

## 7. Discussion and conclusion

When it comes to utilizing electrical stimulation in the nervous system, electrophysiological recordings from an increasing number of neurons and an increasing number of experimental settings are in danger of being corrupted by the infamous stimulation artifact. The currently used gold standard of blanking the recording system during stimulation pulses to make the amplifiers invulnerable against saturation and potential damage may very well introduce unexpected errors in precision analysis and reconstruction. Starting from theoretical considerations, we urge to a very cautious and well-reflected use of blanking as it inadvertently introduces loss of information by shrouding perfectly well-defined spikes relevant for statistics.

Even though it seems evident that the usage of any type of blanking circuit in real spike recordings is to be with great caution and its effects need to be studied prior to use, this report is, to our knowledge, the first one to quantify the detrimental effect blanking may have with respect to the actual event counts at clinically relevant stimulation frequencies. We approached this quantification by the known theoretical description of firing activity based on neuronal interspike-interval histograms.

Blanking nullifies the very histogram bins, which coincide with the blanking window period and width within this distribution. As the histogram can be modeled by a Gamma function, it is possible to estimate the number of spikes affected, depending on the window width T_B_, its frequency F_ST_ and the average firing rate R, of the cell type under consideration. As the Gamma function is not limited in time, we restrict calculations to the 99.95 percentile and have to report on loosing up to 27% of spikes with a quite common blanking window of 2ms and therapeutically relevant 130 Hz stimulation.

In order to validate our theoretical evaluation, we used high SNR data from cell attached recordings of Cholinergic interneurons from the nucleus accumbens of rats. They were collected in context of a critical re-assessment of the effects of HFS in slices (Xie, T. Heida, et al. 2014) and corroborate the theoretical results of substantial spike loss by blanking within the error margins of our automated spike detector. In conclusion, we urge establishing confidence in spike statistics during electrical stimulation electrophysiology by taking independent safeguards like pharmacological controls and careful experimental design.

## 8. Impact statement

Blanking (artifact removal by temporarily grounding input), depending on recording parameters, can lead to prominent spike loss. Utmost caution is advised when using blanking circuits.

## 10. Appendix

To counter electrical stimulation artifacts, several signal processing techniques have been developed over time. These efforts have been made to improve the quality of the signal for further studies later. Different techniques and methods for artifact removal such as Blanking circuits (O’Keeffe et al. 2001), detecting the template of the stimuli artifact (Hashimoto et al. 2002), signal filtering (Zhang et al. 2007) and (Gnadt et al. 2003) have been developed over the years with the effort of scientists.

We have tried to design and implement a novel spike detection method that independently detects neuronal spikes in spite of the presence of contaminating stimulation artifact. Using test data, this algorithm was extensively tested on recording segments without and during stimulation.

Furthermore, we also tested our algorithm on recordings acquired during pharmacological intervention, because such interventions tend to change the characteristics of the sign making it a challenge to ascertain when to apply our detection algorithm. The procedure for the spike detection is described as below:

- Signal windowing with adjustable overlaps

After signal windowing, the signal is further subdivided (5ms width with no overlaps in this study). The features of the spikes can change during the recoding due to pharmacological or electrical intervention. For eg., as is illustrated in the **Figure 8**, Gabazine and Atropine can lead to a significant change in the appearance and features of the spikes. Electrical stimulation artifact is another instance that interferes with the spike detection algorithm.

**Figure 8:**
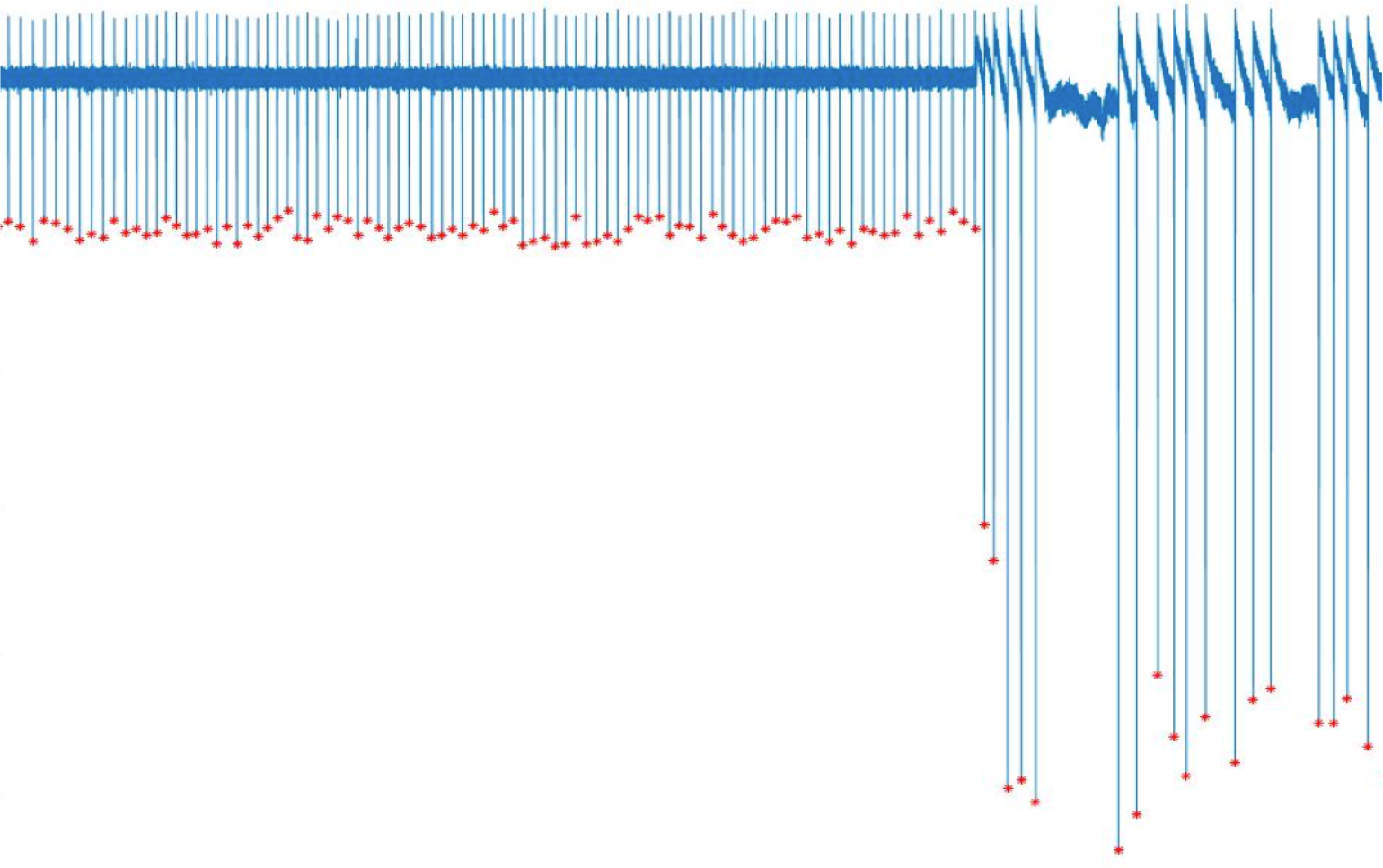
*Spikes’ size can change in an experiment. A change in the amplitude and the spiking rate of a neuron after the Gabazine and Atropin injection. Gabazine blocks GABAa receptors and Atropin inhibits the auto inhibition of the ACH receptors*.

When the stimulation is activated, the signal is buried in a large amount of artifacts, depending on the stimulation frequency and amplitude, as depicted in **Figure 9**.Therefore, the parameters of the spike detection algorithm need to be re-adjusted if there is a change in the pattern.

**Figure 9:**
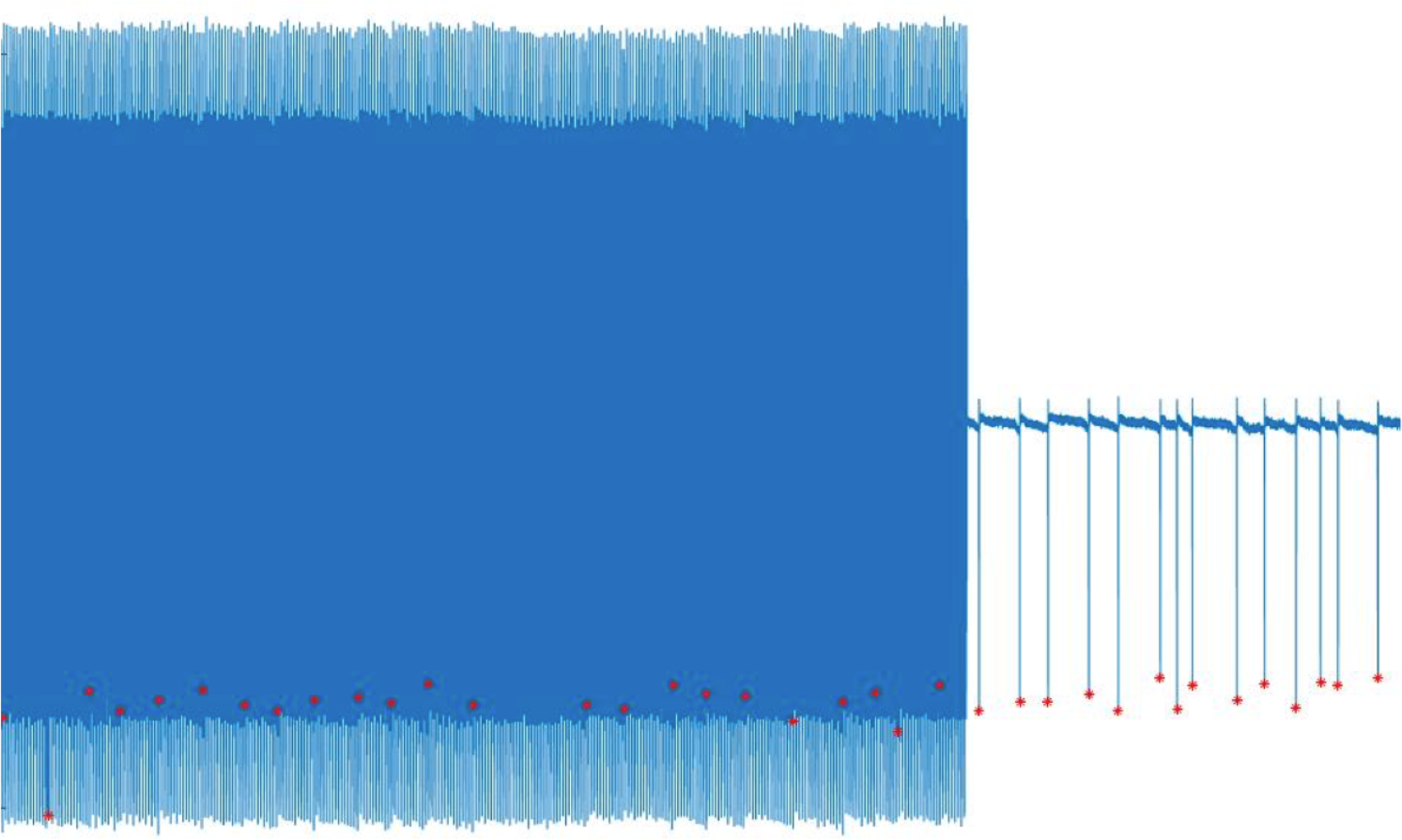
*Electrical stimulation artifact interferes with the signal-processing algorithm*.

- Envelope fitting to the signal

Two envelopes are fitted to the signal in each window, on both the top and the bottom. Based on the experimental setting, a significant change in either of the envelopes may require a new set of parameters in the detection algorithm. The parameters used are valid until the next significant change. It is up to the to the user to define the significance thresholds based on the experimental conditions. **Figure 10** shows an example of the sudden change in the signal due to the cessation of stimulation.

**Figure 10:**
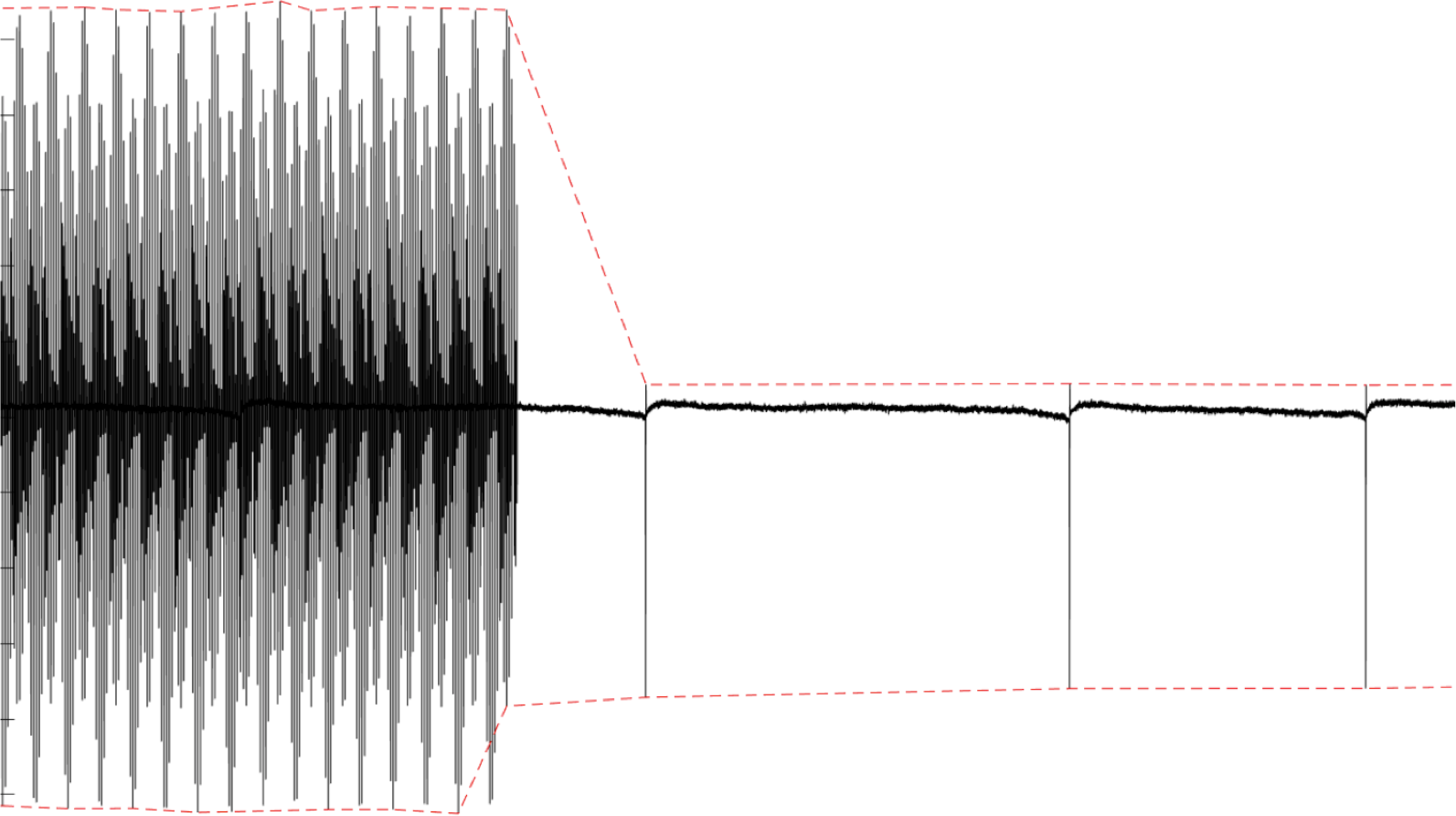
*In each window, two envelopes are fitted to the signal on top and bottom. The user can define the significance of the change, which means the new parameters need to be set*.

- Parameters and Spike detection

The algorithm used in this study looks for the potential spikes using four parameters. Depolarization amplitude (A), rising-falling slopes (B), amplitude of hyperpolarization peaks (C), and distance between the depolarization and hyperpolarization events (D) as shown in the **Figure 11**.

**Figure 11:**
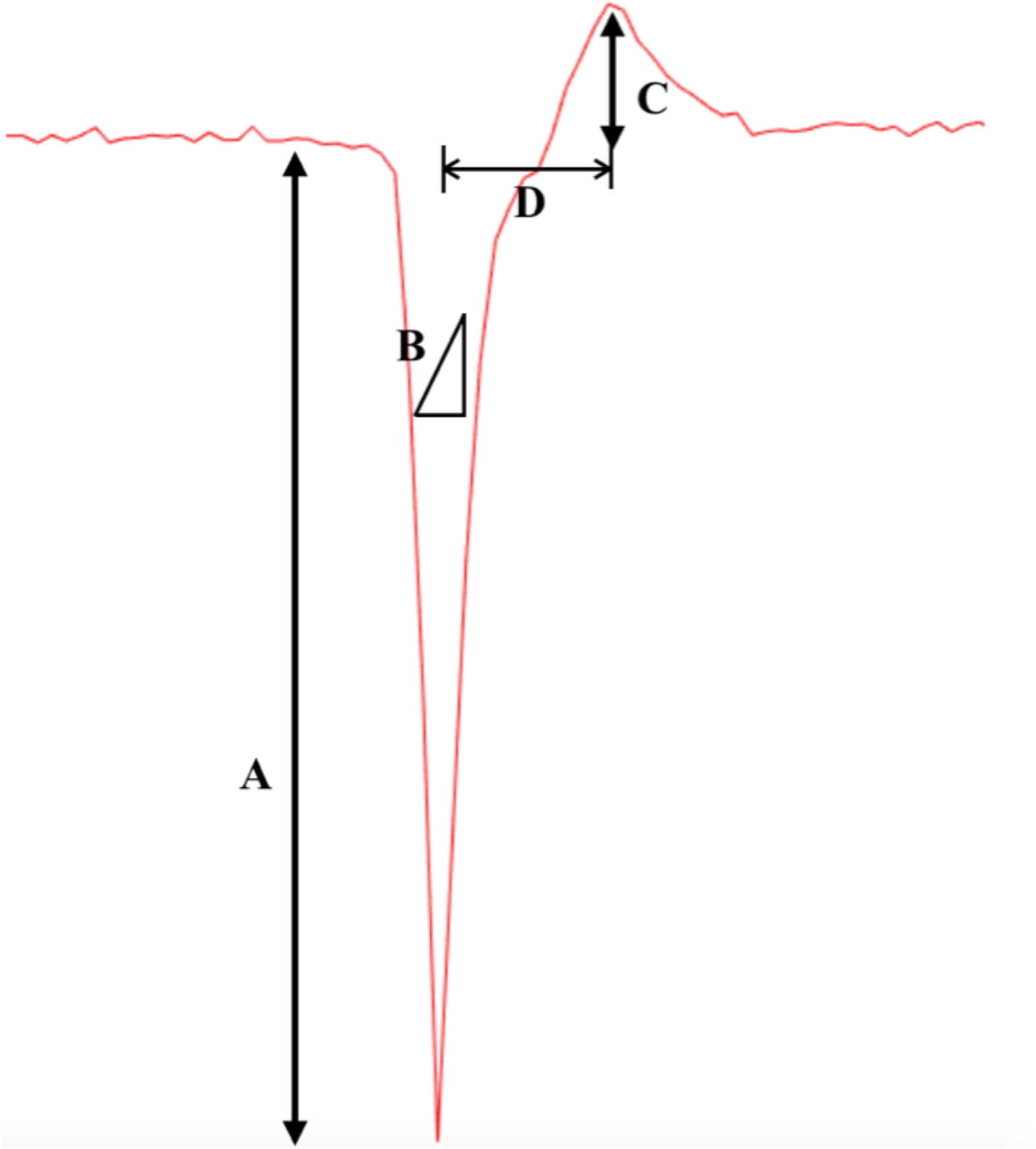
*The parameters used in this study to detect spike are the amplitude (A) and the falling and rising slopes (B) of the depolarization, the amplitude of the hyperpolarization (C), and the time distance between the depolarization and the hyperpolarization event (D)*.

The contamination of the signal with the stimulation artifact can cause a spike overlap in any temporal location and parts of the spike can be lost due to blanking. If three out of four criteria of our algorithm are met, the event is characterized as a spike and saves the amplitude and the time point of the spike. Figure 4 (right) illustrates the possible overlap of electrical stimulation artifacts on a spike, which is chosen from 67 contaminated spikes without blanking. The parameters can be set by averaging over 100 spikes recorded from the target neuron and be used for spike detection. **Figure 12** depicts the flowchart of the method used in this study for spike detection.

**Figure 12:**
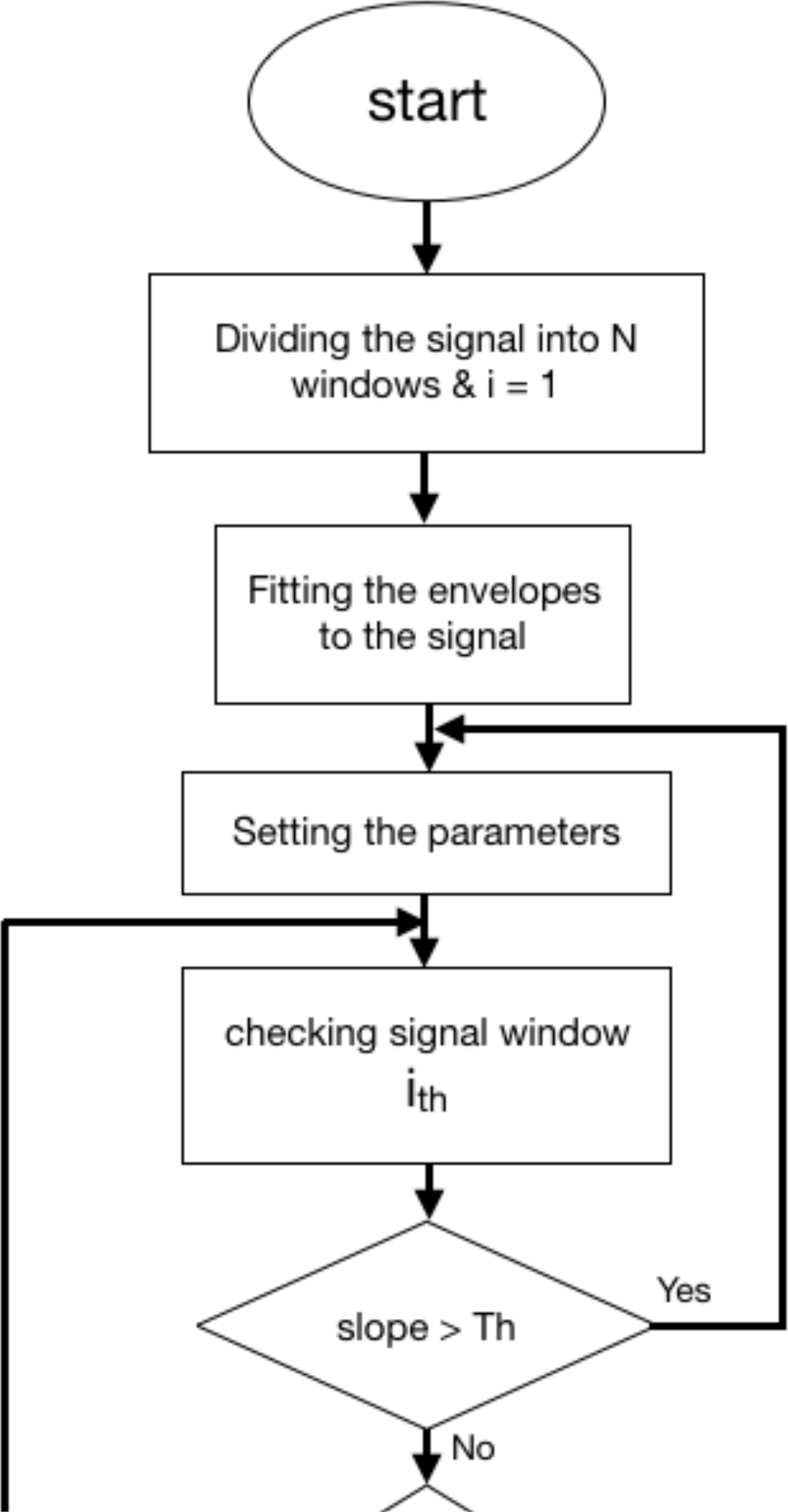
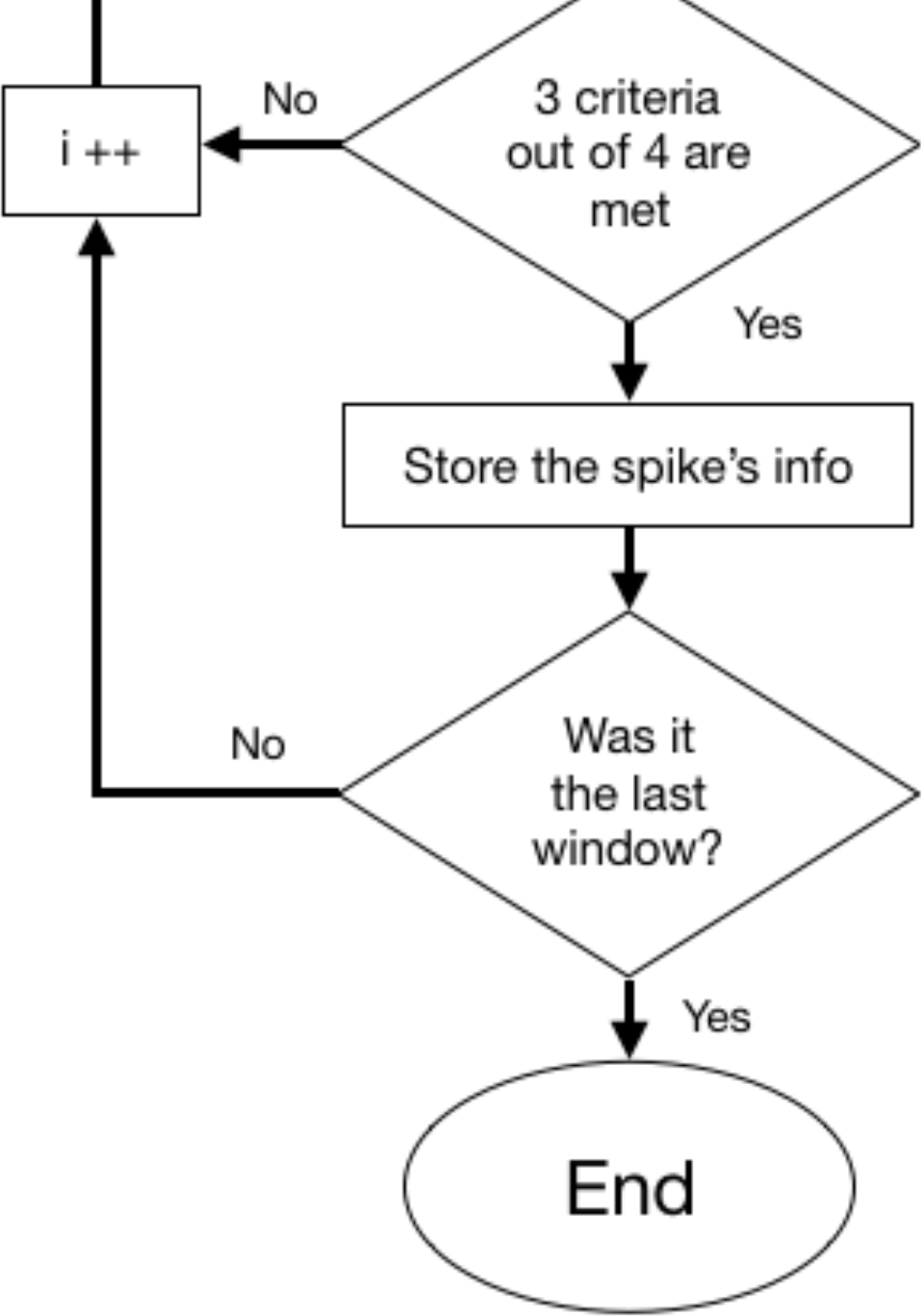
*Flowchart illustrates the method used in this study. According to the expected characteristics of the signal, first the signal should be divided into N windows. Two envelopes are fit to the top and bottom of the signal to quantify trends of the signal. The algorithm looks block by block for the spike and if 3 out 4 criteria which are met, saves the information of that spike otherwise it check the next window.(null)*

